# Xrn1 biochemically associates to the eisosome after the post diauxic shift in yeast

**DOI:** 10.1101/2023.07.19.549689

**Authors:** Baptiste Courtin, Abdelkader Namane, Maïté Gomard, Laura Meyer, Alain Jacquier, Micheline Fromont-Racine

## Abstract

mRNA degradation is one of the main steps of gene expression, and a key player is the 5’-3’ exonuclease Xrn1. In *Saccharomyces cerevisiae*, it was previously shown, by a microscopy approach, that Xrn1 is located to different cellular compartments, depending on physiological state. During exponential growth, Xrn1 is distributed in the cytoplasm, while it is present in the eisosomes after the post-diauxic shift (PDS). Here, we biochemically characterized the Xrn1-associated complexes in different cellular states. We demonstrate that, after PDS, Xrn1 but not the decapping (DCP), nor Lsm1-7/Pat1 complexes, was sequestered in the eisosomes, thus preserving mRNAs from degradation.

## Description

Gene expression is a crucial and well-orchestrated interplay between transcription, translation, and mRNAs and proteins decay. In eukaryotes, cytoplasmic mRNA decay is intimately linked to translation and mediated for a large part by the 5’-3’ mRNA degradation pathway. After mRNA decapping by the DCP complex, with the assistance of its co-factors, the mRNA is degraded by the exonuclease Xrn1 (Parker 2012; Fromont-Racine and Saveanu 2014; for review). This enzyme is highly processive and requires a 5′ monophosphorylated RNA end, produced by decapping (Stevens 1988; Larimer and Stevens 1990). The coupling between mRNA degradation and translation is now well established. First, analyzes of yeast cellular extracts separated on sucrose gradient suggested that the 5’ mRNA degradation could occurs co-translationally (Hu et al. 2009). Second, mapping of the 5’-end by RNA-Seq revealed that degradation intermediates are phased according to a typical codon-specific trinucleotide shift pattern. This observation strongly indicated that at least a fraction of mRNAs degradation occurred during translation (Pelechano et al. 2015). Finally, a cryo-electron microscopy structure of yeast 80S ribosome–Xrn1 nuclease complex showed that Xrn1 can be intimately and specifically bound to the mRNA exit tunnel region of the ribosome, fully consistent with the fact that 5’-3’ mRNA degradation pathway is physically coupled to translation (Tesina et al. 2019).

Gene expression varies, depending on the physiological state of the cells. The *S. cerevisiae* cells must quickly adapt to survive stress conditions. Consequently, a fast reprogramming of gene expression must be set up (DeRisi et al. 1997). Under nutrient starvation, translation of many mRNAs is inhibited (Ashe et al. 2000). This is correlated with Xrn1 cellular localization changes. In response to stress conditions, such as glucose depletion, many mRNA-associated factors involved in mRNA degradation, such as Xrn1 and the DCP complex, are concentrated in processing bodies (P-bodies) (Teixeira and Parker 2007). During the diauxic shift, which happens through the transition from exponential growth to stationary phase, as well as during the post-diauxic shift period (PDS), many cellular modifications occur. The Malinsky’s team previously observed, that Xrn1 is sequestered in the eisosome in PDS, by a microscopy approach (Grousl et al. 2015; Vaškovičová et al. 2017).

The eisosome is a structure associated to the plasma membrane near endocytosis sites. The two main components, Pil1 and Lsp1, share 72% sequence identity and are present in around equimolar proportions. However, despite this strong identity, Pil1 appears to be the major actor of eisosome biogenesis (Walther et al. 2006; Fröhlich et al. 2014). To be efficient, Pil1 and Lsp1 are phosphorylated by two protein kinases, Pkh1 and Pkh2, which are part of the eisosome (Walther et al. 2007). When these two kinases are mutated, the eisosome structure is impaired and endocytosis sites localization is affected suggesting a link between eisosome and endocytosis (Luo et al. 2008). However, the role of the eisosome is far from being understood.

We first identified proteins associated to Xrn1 complex during different physiological states of the cell. We compared the Xrn1-associated complexes obtained by affinity purifications from exponential state extracts (OD_600 nm_ 0.6 and OD_600 nm_ 2), diauxic shift (DS) and Post-diauxic shift (PDS) extracts (Figure 1A). LC-MS-MS identification followed by a quantitative LFQ analysis revealed that the eisosome proteins (red dots), were not enriched with Xrn1-complexes in exponential phase but started to be weakly enriched in DS and were highly enriched with the Xrn1-associated complexes from PDS cells. Among the eisosome proteins, the two main proteins of this structure, Pil1 and Lsp1, were the most abundant and the most enriched with the PDS Xrn1-complexes. In contrast, the DCP complex (Figure 1A, yellow triangles) was less and less present in the Xrn1-associated complexes, from the exponential phase to the PDS, while the presence of the Lsm1-7/Pat1 complex (Figure 1A, yellow dots) was still enriched with Xrn1 (Figure 1A, Dataset 1). A volcano-plot representation revealed the Xrn1-associated complex composition in PDS (Figure 1B, Dataset 1). It highlighted that the most enriched proteins during PDS are the components of the eisosome (red dots). The Lsm1-7/Pat1 proteins (yellow dots) were also enriched but not those of the DCP complex (yellow triangles) nor the ribosome (blue and green dots). A comparison between the Xrn1-TAP associated complexes in exponential phase and PDS (Figure 1C, Dataset 1) distinctly showed that Xrn1 was highly associated to the eisosome in PDS (left part of the graph) and highly associated to the DCP complex and to the ribosome in exponential phase (right part of the graph).

**Figure 1.**
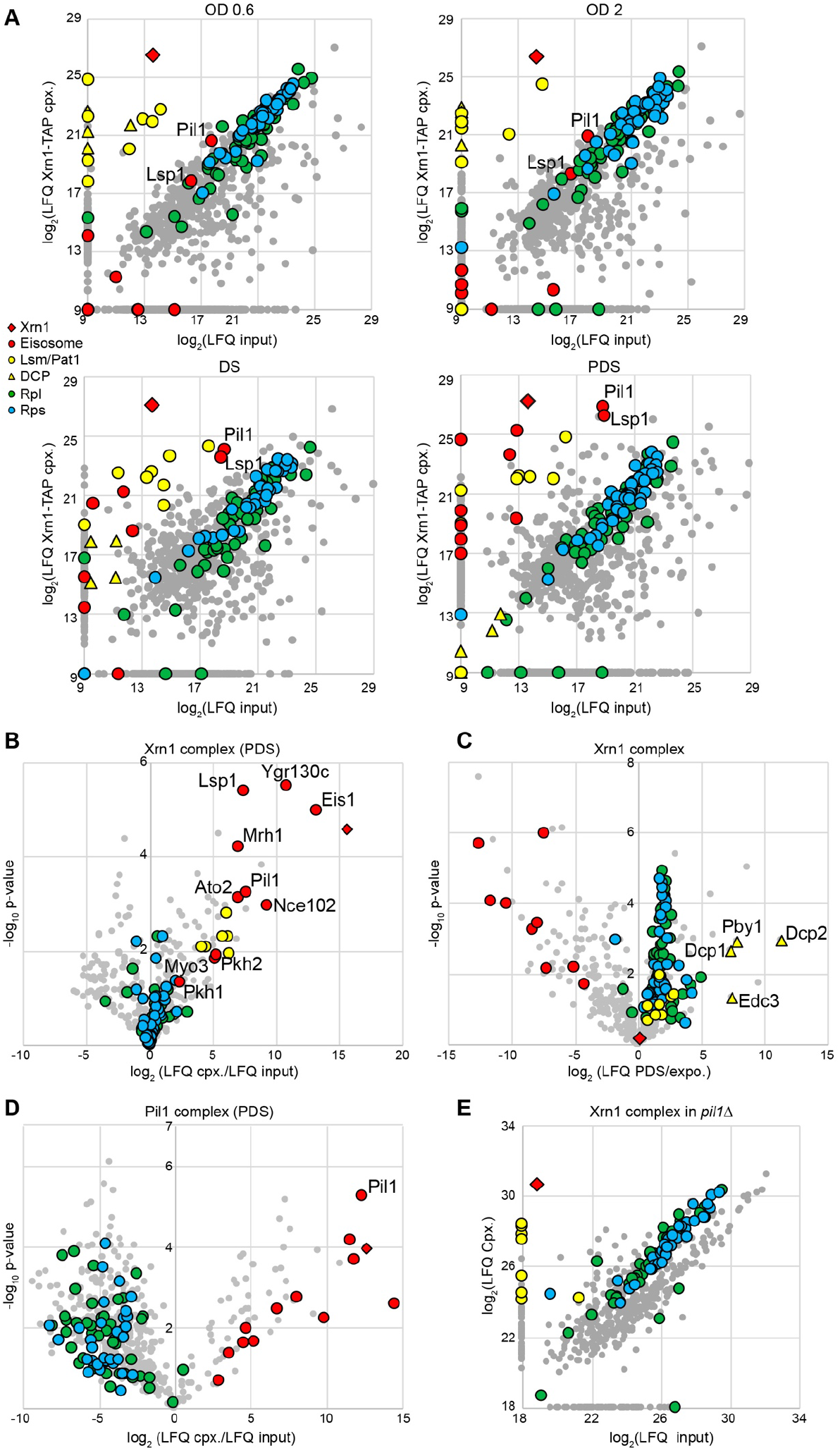
Composition of Xrn1-TAP and Pil1-TAP complexes in different conditions. **A**. Affinity purified Xrn1-TAP associated complexes were subjected to LC-MS/MS identification and quantification. Each panel compares Xrn1-TAP eluate (y-axis) with the total cellular extract (input) (x-axis) from a cell culture in different physiological conditions, as indicated. Each grey dot indicates an identified protein. Red and yellow dots indicate eisosome proteins, and Lsm1-7/Pat1 proteins, respectively, and yellow triangles represent the DCP proteins. Proteins of the large (RPLs) or the small ribosomal subunit (RPSs) are indicated by blue and green dots, respectively. The red diamond designates the Xrn1-TAP bait. **B**. Affinity purification using Xrn1-TAP as bait in PDS condition. Volcano plot shows the fold change (log_2_ LFQ cpx/LFQ input) on the x-axis and the *p*-value distribution (-log_10_ *p*-value) on the y-axis for the proteins identified in the affinity purifications. Colored symbols are as indicated in A. **C**. Comparison of affinity purifications using Xrn1-TAP as bait in exponential and PDS conditions; symbols as in A. **D**. Affinity purifications using Pil1-TAP as bait in PDS condition, as in B. **E**. Scatter plot which compares Xrn1-TAP eluate with the cellular lysate (input) in the absence of Pil1 in PDS; symbols as in A. Purifications for the panel B, C and D were done in triplicate, and in duplicate for the panel E.

To biochemically validate the presence of Xrn1 in the eisosome in PDS and to determine if the Lsm1-7/Pat1 complex is also present in the eisosome, we performed an affinity purification using Pil1-TAP as bait in PDS conditions (Figure 1D, Dataset 1). LC-MS-MS identification followed by a quantitative LFQ analysis indicated that Xrn1 (red diamond) was highly enriched with Pil1 while the Lsm1-7/Pat1 complex was not. In addition, we observed that, in the absence of Pil1, Xrn1 was still associated to the Lsm1-7/Pat1 complex, whereas the eisosome components were missing (Figure 1E, Dataset 1). This is consistent with the observation that in *pil1Δ* mutant strain, the eisosome is completely disassembled (Grousl et al. 2015). In conclusion, we confirmed here, by a biochemical approach, that Xrn1 is localized in the eisosome after post diauxic shift, whereas neither the DCP complex nor the Lsm1-7/Pat1 factors are present in this structure. Our results also suggest that, after PDS and independently of the eisosome, a part of Xrn1 remains associated to the Lsm1-7/Pat1 complex, but not to the DCP complex.

Altogether, these results indicate that, during PDS, Xrn1 is still present but mainly sequestered in the eisosome, probably to protect the mRNAs from full degradation. Nevertheless, a fraction of Xrn1 seems to remain associated to the Lsm1-7/Pat1 complex. It would be of interest to investigate the role of this interaction, independent of the eisosome. This study suggests that sequestration of Xrn1 plays a role in maintaining cell homeostasis during adaptation to nutrient starvation.

## Methods

### Yeast strains and culture conditions

All the strains derive from the BY4741 strain (*MATa, ura3Δ0, his3Δ1, leu2Δ0, met15Δ0*). LMA5015, Xrn1-TAP:HIS3MX6 and LMA5434, Pil1-TAP:HIS3MX6 are from the TAP collection (Ghaemmaghami et al. 2003). LMA5512, Xrn1-TAP:HIS3MX6 *pil1*Δ::KANMX4 was constructed by transformation of the LMA5015 strain with a PCR fragment obtained with *pil1*Δ::KANMX4 strain from the Euroscarf collection using oligonucleotide fw: GAATGGACACTAGACTCTGC and oligonucleotide rv: GGGAACAGAAATGATTATCTGTCC. The cell cultures were grown at 30°C in YPGlu medium, up to an OD_600nm_ 0.6 or 2.0. To collect the DS and PDS samples, the cultures were diluted to an OD_600nm_ 0.005 and grown until they reached the Diauxic State (DS) after 19h of culture or Post Diauxic State (PDS) after 37h of culture. 4000 OD_600nm_ of cell cultures were collected, washed with cold water and stored at -80°C.

### Affinity purification

The cells were resuspended with lysis buffer (20mM Hepes pH7.4, 10mM MgCl_2_, 100mM KOAc) containing a protease-inhibiting reagent (Roche) (1mL per mg of cells). Then, 500 µL of acid-washed glass beads were added per mL of cell suspension and vortexed three-time 40 sec. at 4°C, 6m/sec. (MP FastPrepTM, Fisher Scientific). The suspension was centrifuged for 20 min. at 4°C, 4000 rpm. 0.5% Triton was added to the supernatant, followed by 25 µL of covalently coupled IgG-Dynabeads® magnetic beads resuspended in lysis buffer. The lysate was then incubated for 2 hours at 4°C under agitation. The magnetic beads were recovered and washed five times in washing buffer (20mM Hepes pH7.4, 10mM MgCl_2_, 100mM KOAc, 0.5% Triton), then once in lysis buffer. They were resuspended in 2% SDS, 1X TE and eluted by incubation at 65°C for 15 min. The SDS was removed with HiPPR™ Detergent Removal Resin kit (Thermo Fisher Scientific). Proteins were precipitated by Methanol/Chloroform method (Wessel and Flügge 1984).

### Mass spectrometry analysis

The samples were treated as described in (Defenouillère et al. 2013). Briefly, endoprotease Lys-C and Trypsin digested peptides were analyzed on an LTQ-Orbitrap Velos mass spectrometer (Thermo Fisher Scientific, Bremen). The raw data were analyzed with MaxQuant (version 2.0.3.0). Only proteins identified with at least 2 peptides were selected for further quantification analysis. The abundance of the identified proteins was achieved in the Perseus environment (version 1.6.14). Protein group LFQ intensities were log_2_ transformed. After comparison with a control group, using the Student’s t test statistic, the results were represented as volcano plots or scatter plots.

## Extended data

Dataset 1

## Acknowledgements

We are grateful to the proteomics platform of the Institut Pasteur for the availability of the Orbitrap Velos. We thank Gwenael Badis-Bréard and Cosmin Saveanu for discussions and criticism on the manuscript. We are grateful to Lucia Oreus for providing media and buffers used in this study and Christelle Lenormand for her administrative assistance.

## Funding

This work was supported by the ANR-17-CE11-0049-01, the ANR-17-CE12-0024-02 and the ANR-18-CE11-0003-04 Grants from the Agence Nationale de la Recherche, the Institut Pasteur, and the Centre National de la Recherche Scientifique.

## Author Contributions

Baptiste Courtin: Investigation, formal analysis

Abdelkader Namane: Mass spectrometry investigation, formal analysis

Maïté Gomard: Investigation

Laura Meyer: Investigation

Alain Jacquier: Funding acquisition, Writing -review & editing

Micheline Fromont-Racine: Conceptualization, Funding acquisition, Supervision, Writing -review & editing

## Notes

### Competing Interest Statement

The authors have declared no competing interest.

